# Branched actin constrains endosomal cargo to control sorting and fission

**DOI:** 10.64898/2026.03.10.710749

**Authors:** Devin Frisby, Naava Naslavsky, Steve Caplan

**Author notes:** To whom correspondence and material requests should be addressed: Steve Caplan: ^1^ Dept. of Biochemistry & Molecular Biology, University of Nebraska Medical Center, Omaha, NE, 68198.

## Abstract

At the early endosome, cargos are sorted into subdomains; receptors destined for recycling to the plasma membrane are sorted into tubulovesicular structures that undergo fission and release cargo-laden vesicles that traffic along microtubules. Although branched actin has been implicated in the establishment/maintenance of endosomal membrane subdomains, its role in cargo segregation, fission, and recycling has not been extensively studied. Using inhibitors of formin-and ARP2/3-mediated actin assembly, we show that branched actin, but not linear actin, is required for endosome fission and receptor recycling. To examine the spatial relationship between actin and cargo, we transfected cells with constitutively active RAB5 Q79L to generate enlarged endosomes and demonstrated that internalized transferrin localized to discrete endosomal regions adjacent to branched actin. ARP2/3 inhibition disrupted this organization, resulting in broader cargo distribution on the endosomal membrane and coalescence of degradative and retrieval subdomains. Consistent with impaired endosomal sorting and fission, branched actin inhibition led to cargo accumulation. Our findings identify ARP2/3-mediated branched actin as a key regulator of cargo segregation, subdomain maintenance, and fission at the early endosome.

## INTRODUCTION

Endocytic trafficking refers to the internalization of lipids and integral membrane proteins from the plasma membrane (PM) and their subsequent transport through intracellular compartments. It is critical to cellular function that cargos are trafficked to their appropriate destinations, as higher-order processes, including membrane remodeling, signal transduction, cell migration, and control of cell polarity, are regulated by endocytic trafficking (Caswell & Norman, 2008; Cullen & Steinberg, 2018; Wang *et al*, 2000). Since trafficking is tightly linked to cellular homeostasis, perturbations of these pathways have severe consequences for human health (Yarwood *et al*, 2020). Once internalized, the ability of surface-expressed receptors, such as the β2 adrenergic receptor (β2AR), insulin receptor, epidermal growth factor receptor (EGFR), and the glucagon receptor, to continually transmit signals to the cell interior relies on their recycling back to the PM via endocytic compartments (Gilleron & Zeigerer, 2023; Grant & Donaldson, 2009; Naslavsky & Caplan, 2018; Temkin *et al*, 2011). These endocytic pathways are highly regulated and result in rapid PM turnover; for example, macrophages undergo complete turnover of their surface membrane every 33 minutes (Steinman *et al*, 1976).

Newly endocytosed vesicles undergo homotypic and heterotypic fusion to form the early endosome (EE), which serves as the first destination for cargos internalized from the PM (Jovic *et al*, 2010; Naslavsky & Caplan, 2018). The EE is also referred to as the sorting endosome (SE), as it serves as a sorting station for the recently internalized cargos (Jovic *et al*., 2010). At the EE, receptors are sorted to their respective trafficking pathways (Brown *et al*, 1983; Farquhar, 1983; Steinman *et al*, 1983). Receptors can be sorted to the *trans-*Golgi network (TGN) via retrograde trafficking, to lysosomes for degradation, or to the PM via recycling pathways (Naslavsky & Caplan, 2018). Furthermore, receptors destined for recycling to the PM can be transported by different routes: fast recycling directly from the EE, or via trafficking to the perinuclear endocytic recycling compartment (ERC) before returning to the PM (Naslavsky & Caplan, 2018; Xie *et al*, 2016). To facilitate efficient receptor trafficking, receptors are sorted into appropriate subdomains on the endosome (Derivery *et al*, 2012; Derivery *et al*, 2009; Gomez & Billadeau, 2009; Norris & Grant, 2020; Sonnichsen *et al*, 2000; Temkin *et al*., 2011). Receptors internalized by clathrin-mediated endocytosis (CME) or clathrin-independent endocytosis (CIE) arrive on vesicles that fuse with the same EEs (Naslavsky *et al*, 2003). Interestingly, most CME cargos are recycled from the ERC to the PM through Rab11-positive recycling endosomes (REs), while many CIE cargos enter MICAL-L1-decorated tubular recycling endosomes (TREs) for recycling (Xie *et al*., 2016). This segregation of CME and CIE cargos begins at the EE and is maintained at the ERC (Tian *et al*, 2021; Xie *et al*., 2016). Thus, endosomes must actively sort and maintain receptors within these subdomains.

Since the transferrin receptor (TfR) recycles even after removal of its cytoplasmic domain, it was previously thought that cargo recycling is a passive or “default” process for receptors that were not actively sorted for retrograde trafficking or degradation (Cullen & Steinberg, 2018; Dunn *et al*, 1989; Mayor *et al*, 1993). However, recent studies support the notion that recycling is an active process mediated by cargo tail sorting signals and protein complexes that control recycling (Cullen & Steinberg, 2018; Hsu *et al*, 2012; McNally & Cullen, 2018). While various sorting mechanisms at the endosome have been defined, the molecular details governing these processes remain incompletely understood, especially with regard to mechanisms that maintain receptor segregation after sorting. Cargo sorting into endosomal subdomains at EE is achieved through recognition by protein complexes, the spatiotemporal regulation of phosphoinositide generation, and the actin network. For example, receptors targeted to the lysosome for degradation are ubiquitinated and recognized by the endosomal sorting complex required for transport (ESCRT) complex (Bache *et al*, 2003; Vietri *et al*, 2020). The heterotrimeric complex of VPS26-VPS35-VPS29, referred to as the retromer, cooperates with a heterodimeric combination of SNX1/2-SNX5/6 proteins to sort cargos for retrograde trafficking to the TGN (Arighi *et al*, 2004; Kvainickas *et al*, 2017; Seaman, 2004; Simonetti *et al*, 2017). Further contributing to subdomain organization of endosomes, degradative subdomains, marked by HRS, are distinct from TGN-targeted domains, marked by either SNX1 or Shiga toxin (Popoff *et al*, 2009; Zhang *et al*, 2018). The retromer complex is also involved in regulating receptor recycling through its interaction with SNX27, which binds directly to cytoplasmic motifs on cargos, such as β2AR, to sort them for retromer-mediated recycling (Temkin *et al*., 2011). Retriever is another protein sorting complex, comprised of VPS26C, VPS35L, and VPS29, and it interacts with SNX17 to regulate the recycling of receptors, such as β1-integrin (Bottcher *et al*, 2012; McNally *et al*, 2017). The retromer and retriever complexes localize to the same domain of the endosome, termed the retrieval domain, which is distinct from the HRS-associated degradation domain (McNally *et al*., 2017; Popoff *et al*., 2009). Although endosomal subdomains appear to be organized by the final destination of their cargos, how these domains are established and maintained is not completely understood. Moreover, the precise role of actin in subdomain generation and maintenance remains unclear.

In contrast to the direct specificity of cargo selection, actin has been hypothesized to function primarily as a barrier that prevents unwanted diffusion and maintains segregation at the endosomal membrane. The Wiskott-Aldrich Syndrome protein and scar homologue (WASH) complex is localized to EEs and promotes ARP2/3-mediated branched actin polymerization (Derivery *et al*., 2012; Derivery *et al*., 2009; Gomez & Billadeau, 2009; Puthenveedu *et al*, 2010). The retromer recruits the WASH complex to endosomes via direct interaction, and the WASH complex recruits the retriever through indirect interaction with the COMMD/CCDC22/CCDC93 (CCC) complex (Gomez & Billadeau, 2009; Harbour *et al*, 2012; Jia *et al*, 2012; McNally *et al*., 2017; Seaman *et al*, 2013). The interaction of cargo sorting complexes with an actin nucleation-promoting factor highlights a potential connection between actin polymerization, cargo sorting, and subdomain establishment and maintenance at EE. Indeed, the WASH complex associates with the retromer-and retriever-containing retrieval domains that contain receptors destined for recycling, such as the β2AR (Cullen & Steinberg, 2018; Derivery *et al*., 2012; Gomez & Billadeau, 2009; McNally *et al*., 2017; Puthenveedu *et al*., 2010). However, the role of branched actin in the maintenance of multiple subdomains on endosomes, and the precise impact of branched actin depolymerization on cargo segregation, fission, and recycling has not been extensively studied.

Upon cargo sorting into subdomains, cargos destined for recycling enter tubulovesicular structures that undergo fission; the released cargo-laden vesicles then traffic to the PM along microtubules. The WASH complex promotes ARP2/3-mediated branched actin polymerization at the neck of budding vesicles on endosomes (Gomez & Billadeau, 2009; Jia *et al*., 2012; Seaman *et al*., 2013), which generates mechanical forces required for fission (Derivery *et al*., 2009; Gomez & Billadeau, 2009; Jia *et al*., 2012). Our current model for fission at endosomes is that FCHSD2 promotes WASH-mediated branched actin polymerization to drive tubulation of budding vesicles (Delevoye *et al*, 2016; Derivery *et al*., 2009; Frisby *et al*, 2024; Naslavsky & Caplan, 2023), and Coronin 1C (Hoyer *et al*, 2018; Striepen & Voeltz, 2022) and/or Coronin 2A (Dhawan *et al*, 2022) are recruited to attenuate further actin polymerization, allowing GTPases/ATPases, including Dynamin2 and EHD1, to complete the fission (Deo *et al*, 2018; Duleh & Welch, 2010; Naslavsky & Caplan, 2023; Sharma *et al*, 2009). This model highlights the critical temporal regulation of actin polymerization during endosome fission.

Upstream of fission, actin has been proposed to serve as a physical barrier that inhibits diffusion of cargos across the endosomal membrane to establish specific microdomains on the endosome (Puthenveedu *et al*., 2010; Simonetti & Cullen, 2019). However, the small size of endosomes and the highly dynamic nature of actin have been major barriers in resolving cargo sorting at EE, and the visualization and characterization of distinct endosomal subdomains are complex and understudied. Additionally, it is not well understood how subdomain establishment and cargo sorting are linked to endosome fission. Here, we show that branched actin forms dynamic boundaries that constrain cargo organization on early endosomes, thereby enabling efficient sorting and fission.

## RESULTS

### ARP2/3-mediated branched actin, but not formin-mediated linear filaments, is required for endosome fission

The presence of ARP2/3-mediated branched actin at endosomes has been clearly established (Derivery *et al*., 2012; Derivery *et al*., 2009; Frisby *et al*., 2024; Ohashi *et al*, 2011; Puthenveedu *et al*., 2010). Formins, nucleators of linear actin filaments, have also been implicated in regulating actin polymerization at endosomes, primarily to mediate endosome motility through asymmetric actin polymerization forces (Gasman *et al*, 2003; Wallar *et al*, 2007), but also potentially to regulate receptor recycling (Gong *et al*, 2018). To determine the role of distinct actin nucleation pathways in endosome fission, we used the ARP2/3 inhibitor CK-666 (Nolen *et al*, 2009) and the formin inhibitor SMIFH2 (Rizvi *et al*, 2009). HeLa cells were untreated (Fig. 1A-C), treated with SMIFH2 (Fig. 1D and E), CK-689 (an inactive control for CK-666 (Nolen *et al*., 2009); Fig. 1F and G), or CK-666 (Fig. 1H-J). Cels were then immunostained for endogenous EEA1 to mark early endosomes (Fig. 1A, D, F, H). Cells were also immunostained for cortactin (Fig. 1B, E, G, I), an ARP2/3-binding proteins and a marker of branched actin (Uruno *et al*, 2001; Weed *et al*, 2000). Neither treatment with SMIFH2 nor CK-689 disrupted cortactin localization at endosomes, whereas CK-666 treatment reduced endosome-associated branched actin compared with untreated cells (Fig. 1A-J).

**Figure 1.**
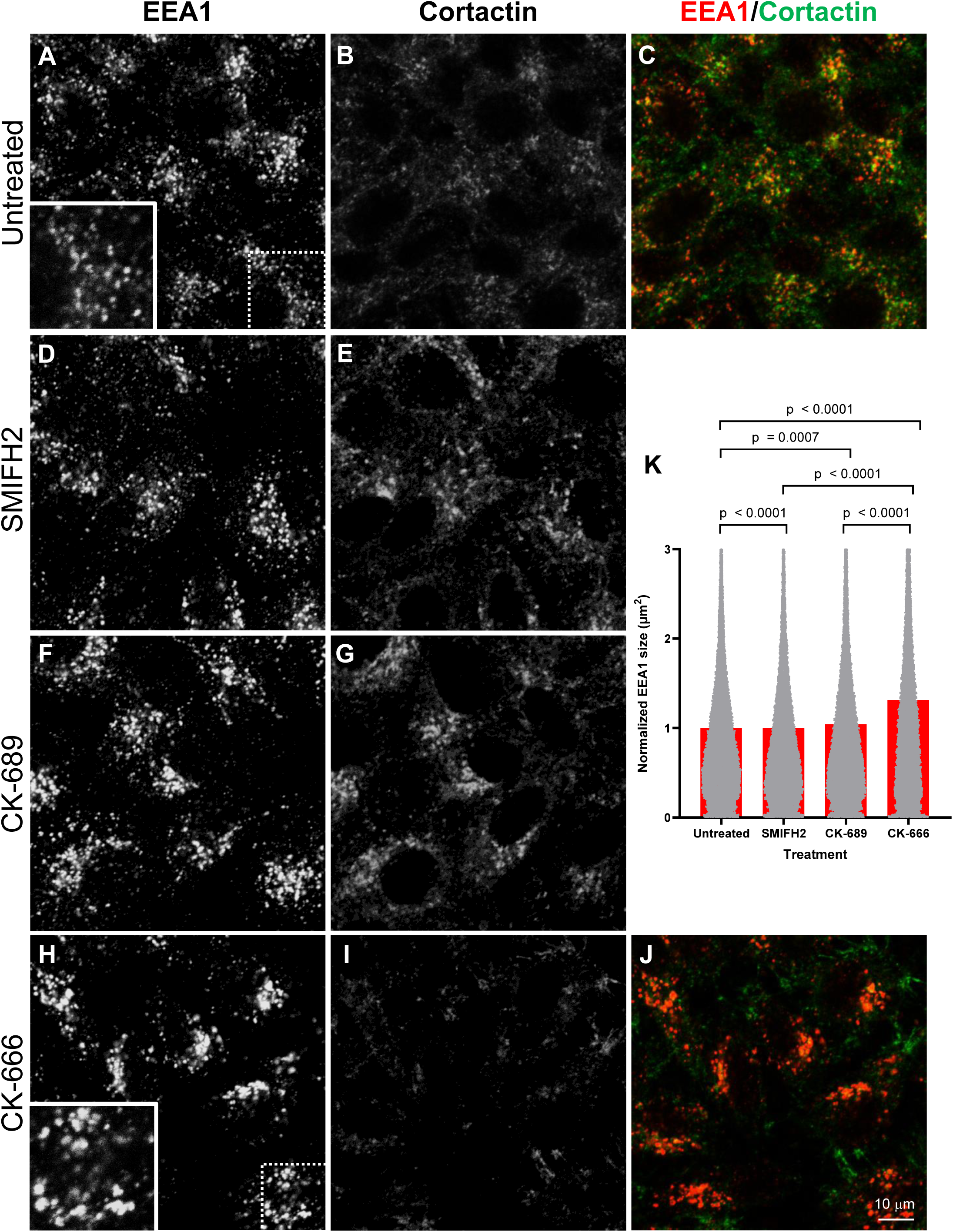
Inhibition of branched actin leads to increased endosome size. **(A-J)**. HeLa cells were (**A-C**) untreated, (**D-E**) treated with the formin inhibitor SMIFH2 (25 µM), (**F-G**) treated with the inactive control CK-689 (300 µM), or (**H-J**) the ARP2/3 branched actin inhibitor CK-666 (300 µM) for 50 min. Cells were fixed and stained with EEA1 and cortactin. Merged images (panels **C** and **J**) show decreased cortactin at endosomes upon CK-666 treatment. **(K)**. Quantification of the effects of the inhibitors on endosome size as depicted in A-J. Imaris software was used to render EEA1-decorated endosomes as surfaces, and endosome size for each treatment (µm^2^) was normalized to the average endosome size of the untreated group. Quantification represents 30 images from three independent experiments, including ∼50,000 endosomes per treatment. A two-tailed Mann-Whitney nonparametric test was used to determine *p*-values between treatment groups.

The WASH complex, a nucleator of ARP2/3-mediated branched actin, is required for fission at endosomes (Derivery *et al*., 2009; Gomez & Billadeau, 2009). Indeed, when we compared endosome size as a surrogate for fission for untreated control cells, SMIFH2-treated cells, CK-689-treated cells, and CK-666-treated cells, only the branched actin inhibitor CK-666 had a major impact on fission, increasing endosome size by ∼30% at 50 min of treatment. Because the dataset included more than 50,000 endosomes per condition, even small differences reached statistical significance. The increase in endosome size with CK-666 is time-dependent, increasing steadily between 15-50 min of treatment (Fig. EV1). A progressive, time-dependent increase in EE size suggests that ARP2/3 inhibition impairs endosome fission. Together, these results demonstrate that endosome fission depends on ARP2/3-mediated branched actin but not formin-mediated linear actin.

### Branched actin regulates receptor recycling to the PM

We next hypothesized that branched actin inhibition at EEs might impair receptor recycling. To address this, HeLa cells were treated with the specified inhibitor and incubated with Tf, a clathrin-dependent cargo, for 10 min to allow internalization. As expected, treatment with SMIFH2, and CK-666 led to decreased Tf internalization (Fig. EV2), consistent with established roles for formins in regulating plasma membrane tension (Djakbarova *et al*, 2021; Scholz *et al*, 2024), and for branched actin in CME (Benesch *et al*, 2005; Lamaze *et al*, 1997; Merrifield *et al*, 2004). Notably the inactive analog CK-689 also reduced internalization, suggesting that this effect may reflect cortical stiffening or other ARP2/3-independent perturbations. Therefore, internalization values were normalized prior to analysis of receptor recycling.

To assess the impact on receptor recycling, cells were incubated with Tf for 10 min of internalization and were chased in complete media in the presence or absence of the inhibitors. Remaining intracellular Tf was measured by determining the fluorescence intensity of the labeled cargo in the cells after 40 min of chase, and the levels were normalized to the uptake values for that treatment (Fig. EV2). Modest delays in Tf recycling were observed for SMIFH2-and CK-689-treated cells (Fig. 2E-G; quantified in Figure 2I). However, treatment with CK-666 showed a more significant recycling defect (Fig. 2H, quantified in Figure 2I), underscoring the role of branched actin in receptor recycling.

**Figure 2.**
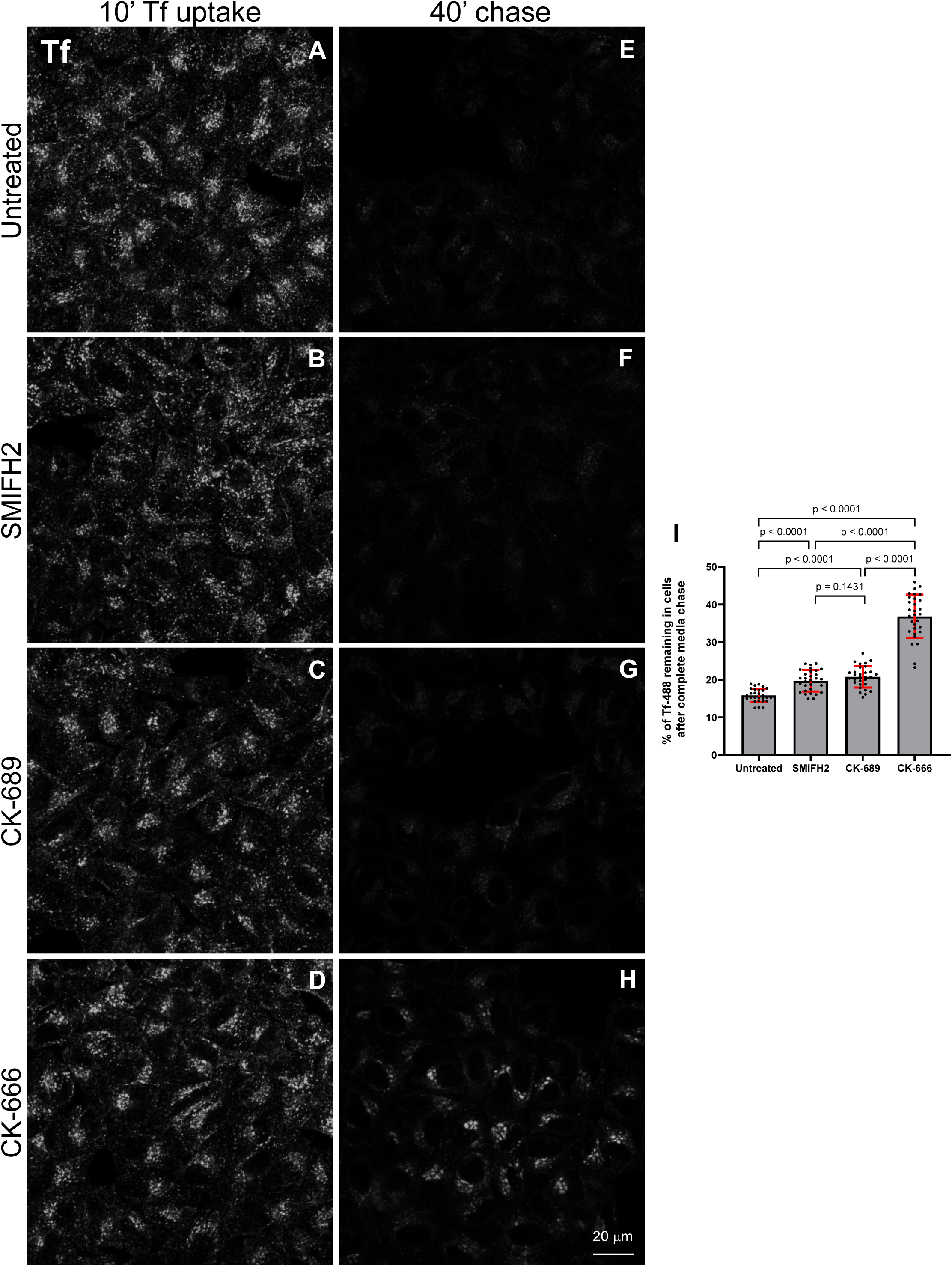
Transferrin recycling is impaired upon ARP2/3 inhibition. **(A-D)**. HeLa cells on coverslips were incubated with fluorophore-labeled transferrin (Tf-488) for 10 minutes of uptake. Cells were (**A**) untreated, (**B**) treated with 25 µM SMIFH2, (**C**) treated with 300 µM CK-689, or (**D**) treated with 300 µM CK-666 during the transferrin uptake. Images are representative of a treated coverslip after uptake. **(E-H)**. Cells were chased in (**E**) complete media, (**F**) complete media containing 25 µM SMIFH2, (**G**) complete media containing 300 µM CK-689, or (**H**) complete media containing 300 µM CK-666 for 40 minutes to allow recycling. Images are representative of cells on an treated coverslip after chase. **(I)**. The arithmetic mean intensity of each image was analyzed using Zeiss Zen Blue software after uptake and chase. For each treatment condition, the internalized mean for the uptake was set at 100%, and the arithmetic mean intensity for each recycling image was expressed as a percentage of the normalized uptake for that condition. Quantification represents 30 images from three independent experiments. Statistical significance was determined using a two-tailed unpaired *t*-test.

Based on the impaired fission observed with CK-666 treatment, we hypothesized that the delayed receptor recycling was driven by the failure of Tf-containing vesicles to be released from endosomes. To test this and determine if Tf is maintained at the EE membrane, we measured the amount of Tf retained at EE after 15, 30 and 45 min of chase (Fig. 3E). As demonstrated at the 30 min time point, significantly more Tf is retained in EEs upon CK-666 treatment (compare Figure 3C and D to A and B; quantified in F). Indeed, over time, significantly more Tf remains at the EE in cells treated with CK-666, further supporting the role of branched actin in endosomal fission and recycling (Figure 3E).

**Figure 3.**
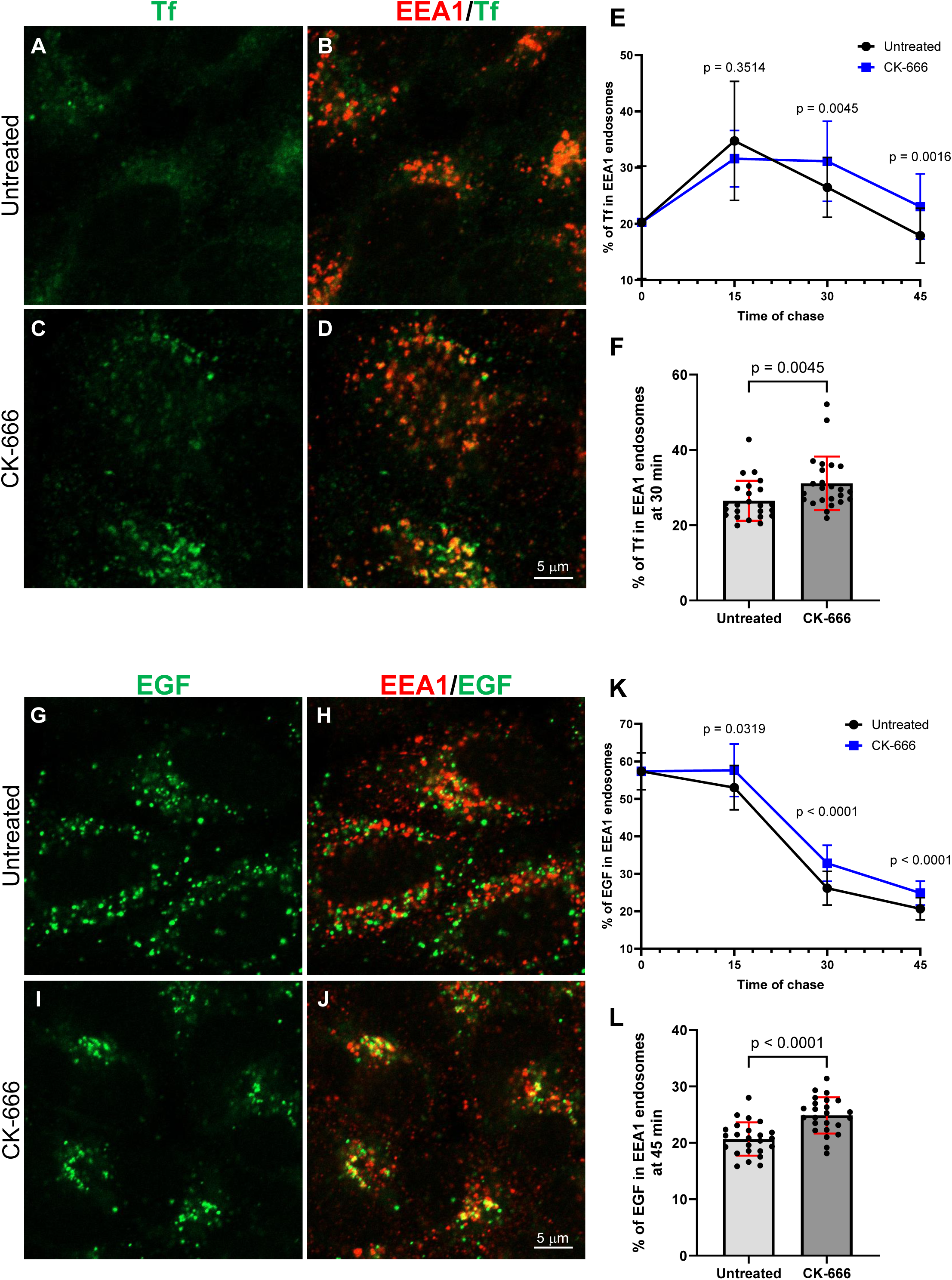
Transferrin and EGF are delayed at early endosomes upon ARP2/3 inhibition. **(A-D)**. HeLa cells were incubated with Tf-488 diluted in complete media for 10 min. Following uptake, cells were chased in (**A-B**) complete media or (**C-D**) in media containing 300 µM CK-666. Representative images are from the 30 min chase time point. **(E).** The percentage of Tf-488 fluorescence in EEA1 endosomes was calculated by measuring the area of Tf-EEA1 overlap as a percentage of total Tf fluorescence area. Statistical analysis was performed between the untreated and CK-666-treated groups at each time point. Quantification represents 24 images from three independent experiments. Statistical significance for the 30 min and 45 min time points was determined using a two-tailed Mann-Whitney nonparametric test, and an unpaired two-tailed *t*-test was used for the 15 min time point. **(F).** A bar graph showing the individual data points from the 30 min time point in (**E**). **(G-J)**. HeLa cells were serum starved for 1 h, then incubated with EGF-488 diluted in complete media for 10 min. Following uptake, cells were chased in (**G-H**) complete media or (**I-J**) in complete media containing 300 µM CK-666. Representative images are from the 45 min time point of media chase. **(K).** The percentage of EGF-488 fluorescence in EEA1-marked endosomes was calculated by measuring the area of EGF-EEA1 overlap as a percentage of total EGF fluorescence area. Statistical analysis was performed between the untreated and CK-666-treated groups at each time point. Quantification represents 24 images from three independent experiments. Statistical significance for the 15 min and 45 min time points was determined using a two-tailed Mann-Whitney nonparametric test, and an unpaired two-tailed *t*-test was used for the 30 min time point. **(L).** A bar graph showing the individual data points from the 45 min time point in (**K**).

Whereas Tf is a cargo that remains bound to the TfR and is efficiently recycled to the PM, EGF-bound EGFR is typically targeted to lysosomes for degradation (Beguinot *et al*, 1984). To assess whether inhibition of the branched actin network at EEs affects EGFR trafficking into lysosomes, we followed the intracellular distribution of fluorescently labeled EGF after internalization and chase. We found that upon CK-666 treatment, there was a similar increase in EGF and its receptor retained in EEA1-containing EEs (Fig. 3, compare I and J to G and H; quantified at 45 min chase in L). Over 15-45 min of chase, the EGF is consistently maintained in EEA1-containing EEs when branched actin is disrupted with CK-666 (Fig. 3L). Overall, branched actin is required for the exit of both recycling and degradation cargo from the EE, consistent with the observation that WASH knockdown impairs EGF entry into lysosomes (Duleh & Welch, 2010).

### Recycling cargos are bound by branched actin on endosomes

The WASH complex is recruited to endosomes via the retromer and also indirectly interacts with the retriever complex (Gomez & Billadeau, 2009; Harbour *et al*., 2012; Jia *et al*., 2012; McNally *et al*., 2017; Seaman *et al*., 2013). This raises the possibility of a role for branched actin not only in terminal EE fission, but also in earlier cargo-sorting steps. However, how the endosomal actin network established during cargo sorting is coupled to endosomal fission remains incompletely understood. The highly dynamic nature of actin at endosomes and their limited surface area complicate the study of EE subdomain establishment. To overcome this, we transfected cells with the GTP-locked RAB5 Q79L mutant (Stenmark *et al*, 1994). Transfection of this mutant caused the formation of enlarged endosomes, which were positive for phalloidin-marked actin and cortactin-marked branched actin (Fig. 4 A-D; quantified in Q). When the RAB5 Q79L-transfected HeLa cells were incubated with the formin inhibitor SMIFH2 or the inactive CK-689, the endosomal actin remained intact (Fig. 4 E-L; quantified in Q). However, as we have previously shown (Frisby *et al*., 2024), when cells were treated with the ARP2/3 inhibitor CK-666, there was a significant decrease in both filamentous actin and cortactin at endosomes (Fig. 4 M-P; quantified in Q). These data are consistent with a role for ARP2/3 in generating branched actin at endosomes.

**Figure 4.**
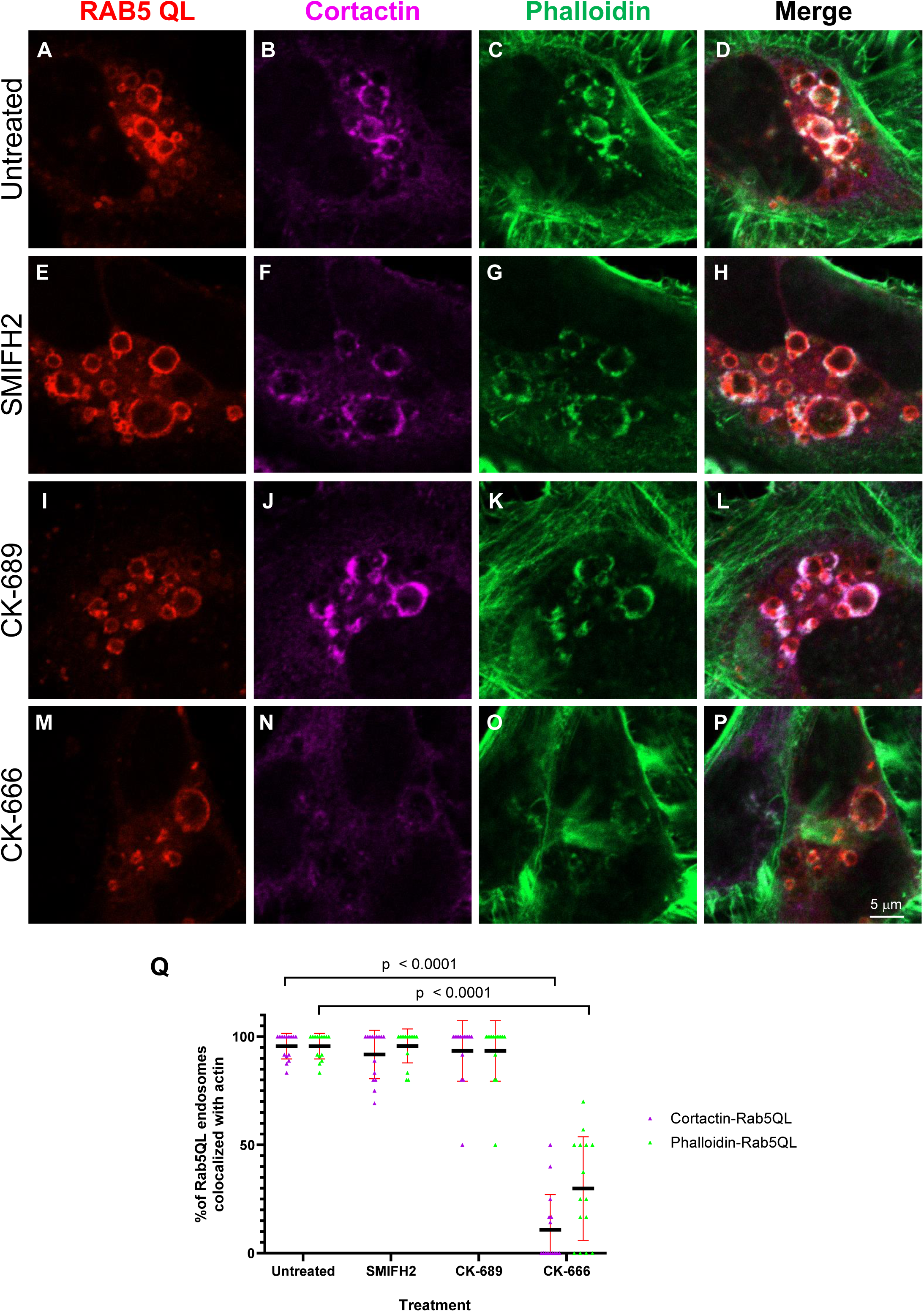
Branched actin inhibition, but not formin inhibition, decreases actin at RAB5 QL endosomes. **(A-P)**. HeLa cells were transfected with mCherry-RAB5 Q79L and were either (**A-D**) untreated or treated with (**E-H**) 25 µM SMIFH2, (**I-L**) 300 µM CK-689, or (**M-P**) 300 µM CK-666 for 20 min. Cells were fixed and co-stained with cortactin and phalloidin to visualize the actin network. **(Q)**. Quantification of (**A-P**). RAB5 Q79L endosomes that colocalized with phalloidin or cortactin were counted as a percentage of total RAB5 Q79L endosomes. Quantification represents 15 images from three independent experiments. A two-tailed Mann-Whitney nonparametric test was used to determine significance between treatment groups. Data comparisons without error bars are not significant (*p* > 0.05).

To visualize actin localization at EE together with internalized receptors, we transfected cells with the RAB5 Q79L mutant, incubated them with Tf, and immunostained cells for cortactin to mark the branched actin around the endosome (Fig. 5 A-L). Regions containing Tf with overlapping or adjacent regions of cortactin were observed (Fig. 5 A-C). To quantify the relationship between cargo and actin distribution, a circular region of interest was plotted around the endosomal membrane that divides the endosome into 360 degrees, and software was designed and applied to measure the fluorescence intensities of Tf and cortactin surrounding the endosome at each individual degree. The generated intensity profiles of the circular ROIs were normalized to their mean fluorescence values, and the number of Tf regions bounded by cortactin was quantified. A region of Tf was designated as “bounded” by cortactin if a Tf peak exceeding 130% of the mean fluorescence was within 20 degrees of a cortactin peak exceeding 130% (Fig. 5 M-P). In untreated cells, Tf was localized to distinct regions, with 63% of Tf regions bounded by cortactin (Fig. 5Q). Treatment with SMIFH2 or CK-689 had little effect on the spatial distribution of actin and Tf on endosomes (Fig. 5Q). In contrast, CK-666 treatment reduced branched actin at endosomes and decreased the fraction of Tf-containing domains bounded by actin (Fig. 5Q). Although branched actin was markedly diminished (compare cortactin in Figure 5K to 5B, E, and H), it was not completely abolished. Accordingly, the proportion of Tf peaks bounded by cortactin decreased to ∼40% in CK-666-treated cells but was not eliminated. Together, these data indicate that Tf, a recycling cargo, occupies discrete endosomal regions that are bounded by branched actin (modeled in Figure 5R,S).

**Figure 5.**
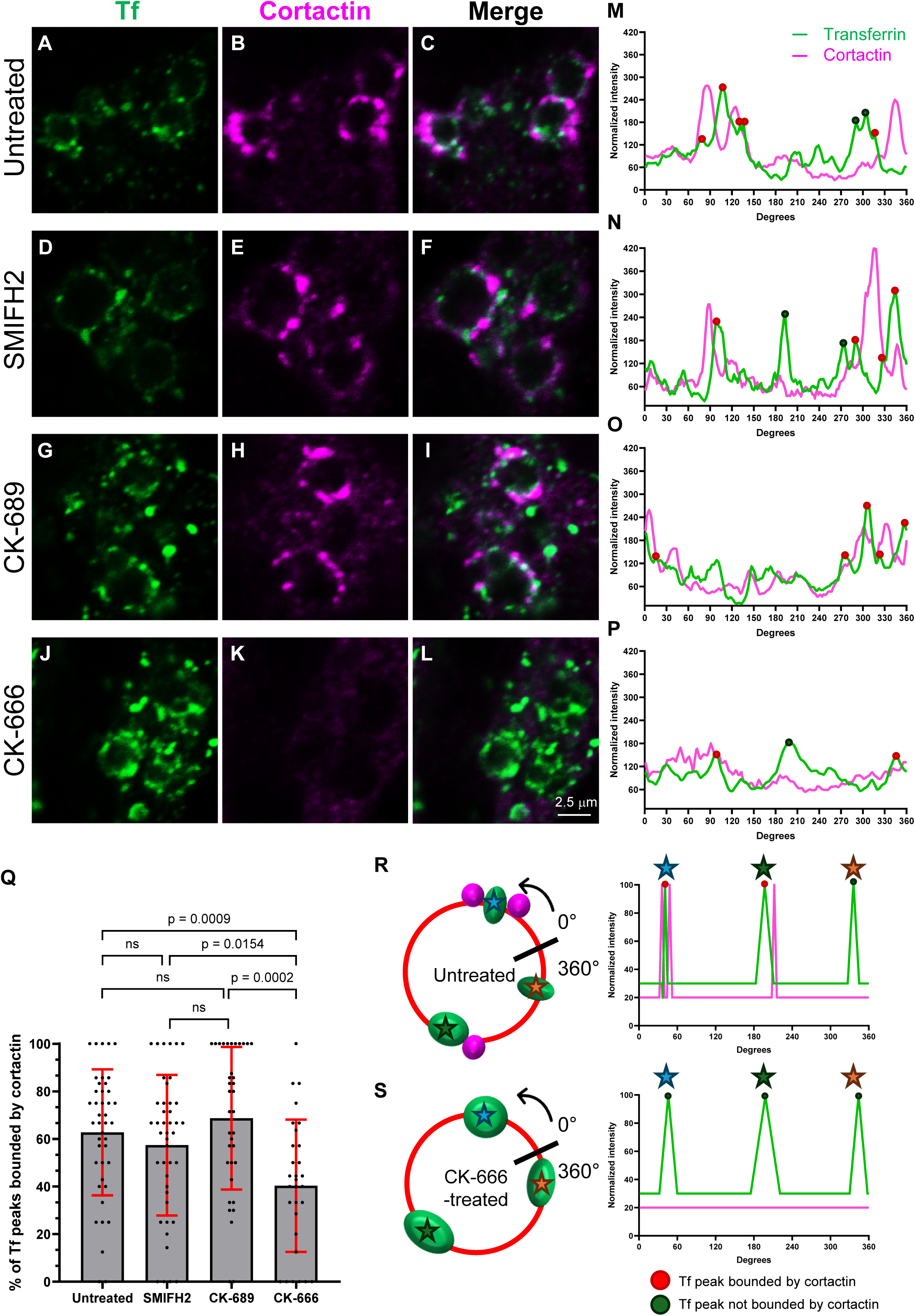
Transferrin is bounded by cortactin at endosomes. **(A-L)**. HeLa cells were transfected with mCherry-RAB5 Q79L and were incubated with (**A-C**) no inhibitor, (**D-F**) 25 µM SMIFH2, (**G-I**) 300 µM CK-689, or (**J-L**) 300 µM CK-666 for 4 min. Following the pre-treatment, cells were incubated with Tf-488 (and their respective inhibitor) for 6 min of uptake. **(M-P)**. Representative fluorescence intensity profiles of endosomes from (**M**) untreated, (**N**) SMIFH2-treated cells, (**O**) CK-689-treated cells, or (**P**) CK-666-treated cells. The intensity profile of Tf is in green, and the intensity profile of cortactin is in magenta. Tf vertices above 130% that occur within 20 degrees of a cortactin value above 130% are represented with a red circle; these peaks are “bounded”. Tf vertices above 130% that are not within 20 degrees of a cortactin value above 130% are represented with a grey circle and are “not bounded”. **(Q).** The percentage of “bounded” Tf peaks was quantified from fluorescence intensity profiles (as represented in (**M-P**)). Quantification is from three independent experiments, and from 37 endosomes for the untreated group, 43 endosomes for SMIFH2-treated, 38 endosomes for CK-689-treated, and 37 endosomes from the CK-666-treated group. Statistical significance was determined using a two-tailed unpaired *t*-test (ns: *p* > 0.05). **(R, S)**. Representative models for the quantification and results of the experiment. A circle was drawn around the endosome membrane (red) that intersects with the regions of Tf (green) and cortactin (magenta), and fluorescence intensity at each degree around the circle was measured. (**R**) Untreated cells have Tf in confined regions on the endosome and are adjacent to regions of cortactin ∼60% of the time. (**S**) CK-666-treated cells have regions of Tf that are broader and “not bounded” by cortactin.

### Branched actin maintains cargo localization to discrete endosomal subdomains

To quantitatively assess the extent to which branched actin serves as a diffusion barrier for membrane bound cargo such as Tf along the endosome membrane, HeLa cells transfected with RAB5 Q79L were incubated with Tf. Fluorescence intensity profiles of Tf around the endosomes were then measured as described above (Fig. 6A-Q). Tf-containing domains were identified from the intensity plots (Fig. 6N–Q), and fluorescence intensities within ±20 degrees of each domain maximum were defined as a Tf “peak”. Domain intensities were normalized to the maximum value within each Tf peak, generating normalized domain profiles that were averaged across endosomes for each treatment group (Fig. 6R). Untreated (Fig. 6A-C; quantified in N), SMIFH2-treated (Fig. 6D-F; quantified in O), and CK-689-treated cells (Fig. 6G-I: quantified in P) exhibited discrete Tf-containing domains on the endosomes (yellow arrows). In contrast, CK-666 treatment disrupted this organization, resulting in significantly broader Tf-enriched regions (Fig. 6J-L; quantified in R, blue arrows). These results support a role for branched actin in maintaining Tf within discrete endosomal membrane subdomains.

**Figure 6.**
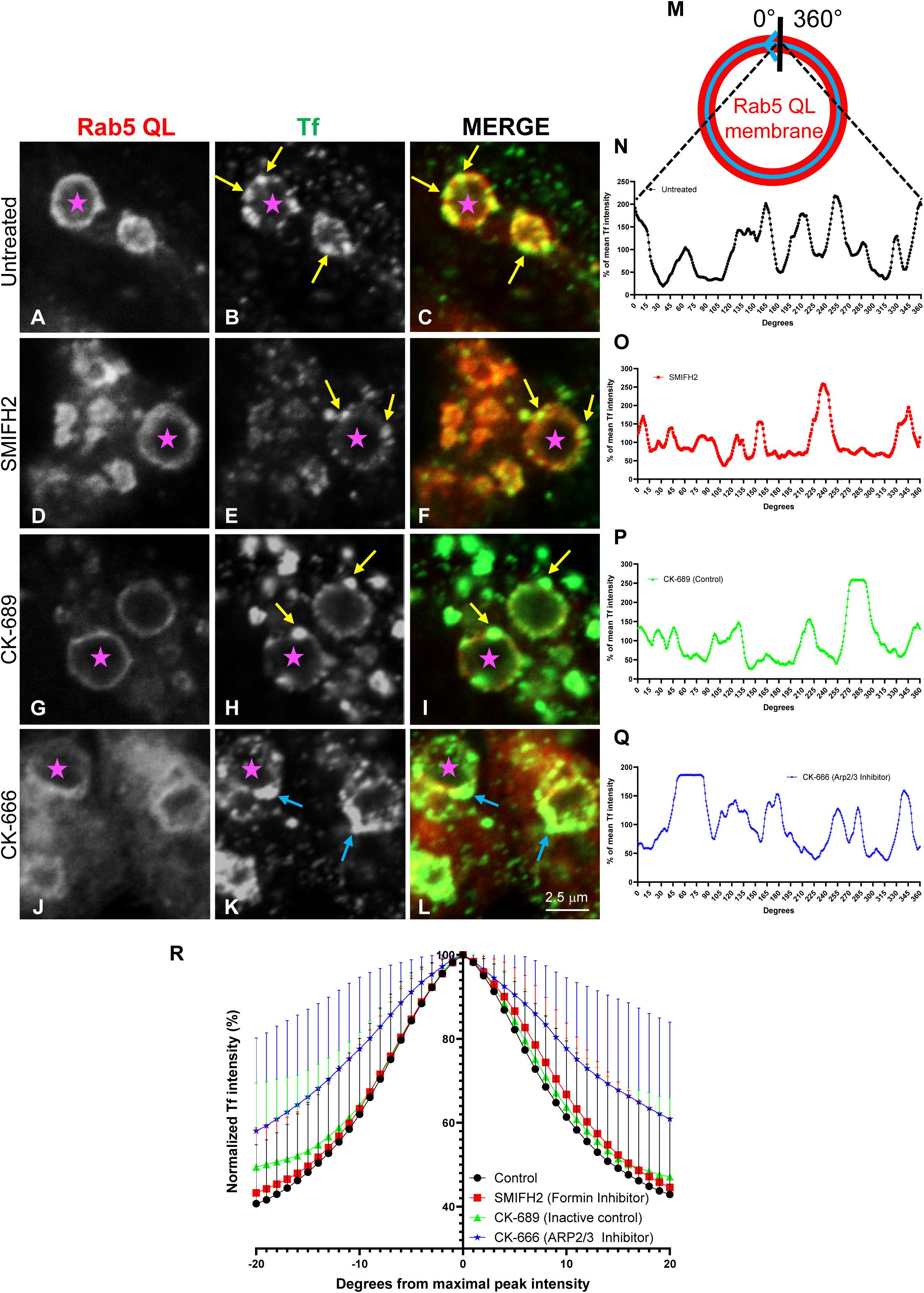
Tf occupies less discrete regions on the endosome when branched actin is inhibited. **(A-L)**. HeLa cells were transfected with mCherry-RAB5 Q79L and were incubated with (**A-C**) no inhibitor, (**D-F**) 25 µM SMIFH2, (**G-I**) 300 µM CK-689, or (**J-L**) 300 µM CK-666 for 4 min. Following the pre-treatment, cells were incubated with Tf-488 (and their respective inhibitor) for 6 min of uptake. Untreated, SMIFH2-treated, and CK-689-treated cells show Tf localized to discrete regions on the endosome (yellow arrows). CK-666-treated cells have broader regions of Tf on the endosome (blue arrows). **(M).** Model for quantification. A circle (blue) was drawn around the endosome membrane (red) that intersects with the regions of Tf, and the fluorescence intensity at each degree around the circle was measured. **(N-Q)**. Representative fluorescence intensity profiles of Tf on the endosome from (**N**) untreated, (**O**) SMIFH2-treated, (**P**) CK-689-treated, and (**Q**) CK-666-treated cells. The profiles depicted are from the endosomes indicated with a magenta-colored star (**A-L**). **(R).** Tf-containing regions from the fluorescence intensity profiles (**N-Q**) were identified, and the fluorescence values ± 20 degrees from the maximum were normalized. These normalized profiles of Tf-containing regions are Tf “peaks”. The graph shows the average profile of 72 Tf peaks from 37 endosomes for the control group, 70 peaks from 43 endosomes from SMIFH2-treated cells, 73 peaks from 38 endosomes from CK-689-treated cells, and 70 peaks from 37 endosomes from CK-666-treated cells. Data are from three independent experiments. The average profile of Tf peaks in the CK-666-treated cells is broader (less discrete) than the other treatment groups. See table in Figure EV3 for statistical information.

Next, we examined whether the branched actin network also restricts receptor diffusion within discrete endosomal subdomains that host cargo destined for degradation, such as the EGF receptor. RAB5 Q79L-transfected cells were incubated with EGF, and the boundaries of the EGF-containing domains were quantified as described above for the Tf-containing domains (Fig. 7). As demonstrated, EGF was confined to discrete domains on the endosomes in untreated, SMIFH2-treated, and CK-689-treated cells (Fig. 7A-I, intensity profiles shown in M-O; quantified in 7Q, yellow arrows). In contrast, CK-666 treatment disrupted this organization, resulting in broader and more diffuse EGF-enriched regions (Fig. 7J-L, blue arrows, intensity profile shown in P; quantified in 7Q). Together, these data indicate that branched actin confines both recycling and degradative cargo within discrete subdomains on the endosome.

**Figure 7.**
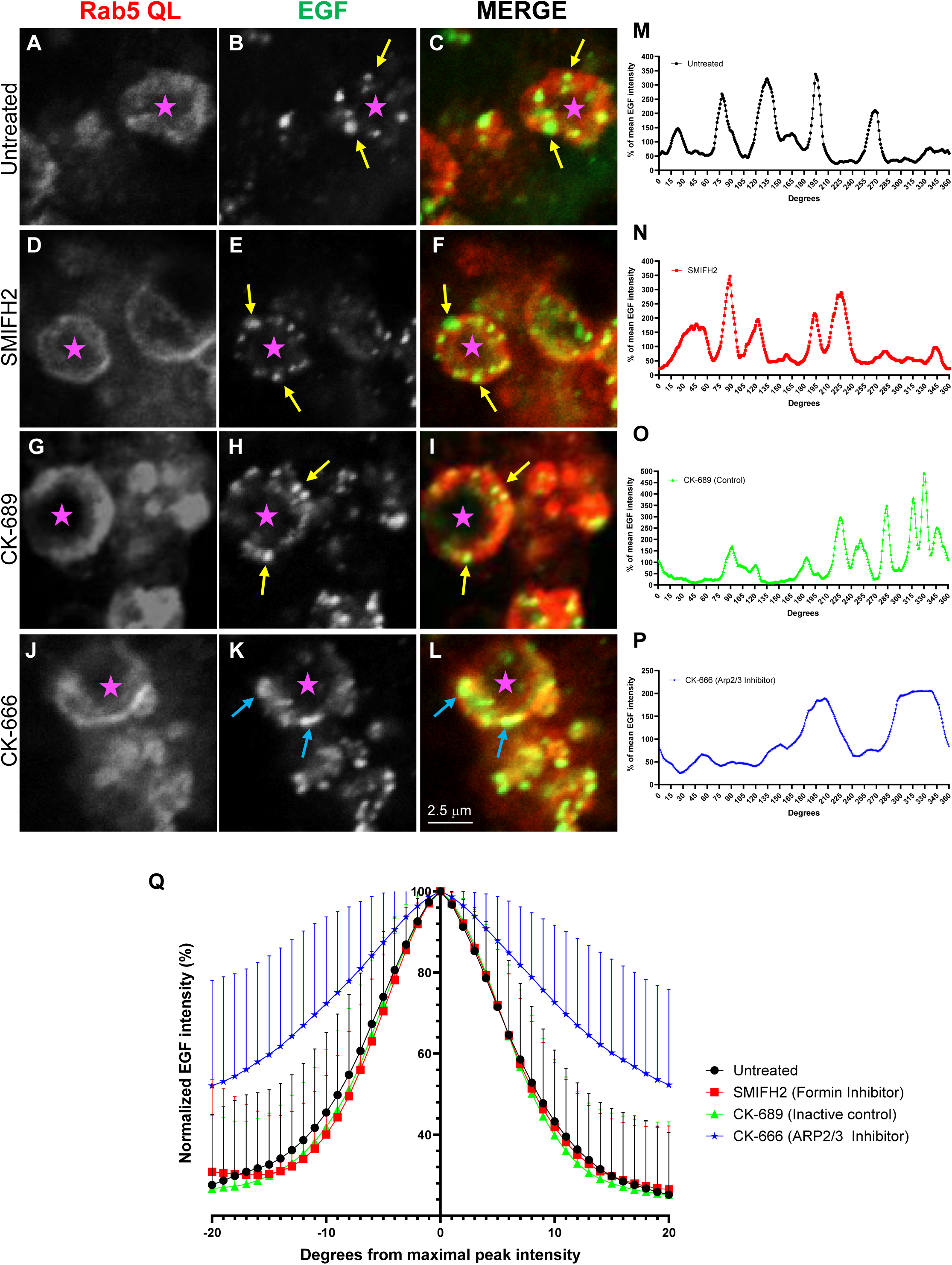
EGF segregation on the endosome is affected by branched actin inhibition. **(A-L)**. HeLa cells were transfected with mCherry-RAB5 Q79L and were incubated with (**A-C**) no inhibitor, (**D-F**) SMIFH2, (**G-I**) CK-689, or (**J-L**) CK-666 for 4 min. Following the pre-treatment, cells were incubated with EGF-488 (and the respective inhibitor) for 17 min of uptake. Untreated, SMIFH2-treated, and CK-689-treated cells show EGF localized to discrete regions on the endosome (yellow arrows). CK-666-treated cells have broader regions of EGF on the endosome (blue arrows). **(M-P)**. Representative fluorescence intensity profiles of EGF on the endosome from the (**M**) untreated, (**N**) SMIFH2-treated, (**O**) CK-689-treated, and (**P**) CK-666 treated cells. The profiles depicted are from the endosomes indicated with a magenta star in (**A-L**). **(Q)**. EGF-containing regions from the fluorescence intensity profiles (**M-P**) were identified, and the fluorescence values ± 20 degrees from the maximum were normalized. These normalized profiles of EGF-containing regions are EGF “peaks”. The graph shows the average profile of 84 EGF peaks from 47 endosomes that were analyzed for the control group, 86 peaks from 51 endosomes from SMIFH2-treated cells, 72 peaks from 39 endosomes from CK-689-treated cells, and 86 peaks from 49 endosomes from CK-666-treated cells. Data are from three independent experiments. The average profile of EGF peaks in the CK-666-treated cells is broader (less discrete) than the other treatment groups. See table in Figure EV4 for statistical information.

### Branched actin maintains distinct retrieval and degradative domains on endosomes

Cargo internalized through clathrin-dependent and-independent endocytosis accumulates on shared endosomal membranes but remains segregated within distinct subdomains (Naslavsky *et al*., 2003). Moreover, degradative and recycling domains occupy spatially distinct regions on the endosome (MacDonald *et al*, 2018; McNally *et al*., 2017; Popoff *et al*., 2009). We therefore asked whether branched actin network is required to maintain segregation between clathrin-dependent and-independent cargo. To this end, HeLa cells were transfected with RAB5 Q79L and simultaneously incubated with antibodies to CD59 (a CIE cargo that undergoes recycling (Cai *et al*, 2011; Cai *et al*, 2014)) and EGF (an EGFR ligand internalized via CME and trafficked to lysosomes (Beguinot *et al*., 1984)). In untreated cells, CD59 and EGF localized to distinct subdomains on the endosomal membrane (Fig. 8A-C). In contrast, inhibition of branched actin polymerization with CK-666 caused EGF-and CD59-containing domains to coalesce, resulting in increased overlap of the cargos on endosomes (Fig. 8D-F; see white arrows). Colocalization was quantified by both Pearson’s and Manders’ colocalization coefficients (Fig. 8 G-I), revealing a significant increase in cargo overlap upon CK-666 treatment. These data indicate that branched actin is required to maintain segregation of recycling and degradative subdomains on the endosome.

**Figure 8.**
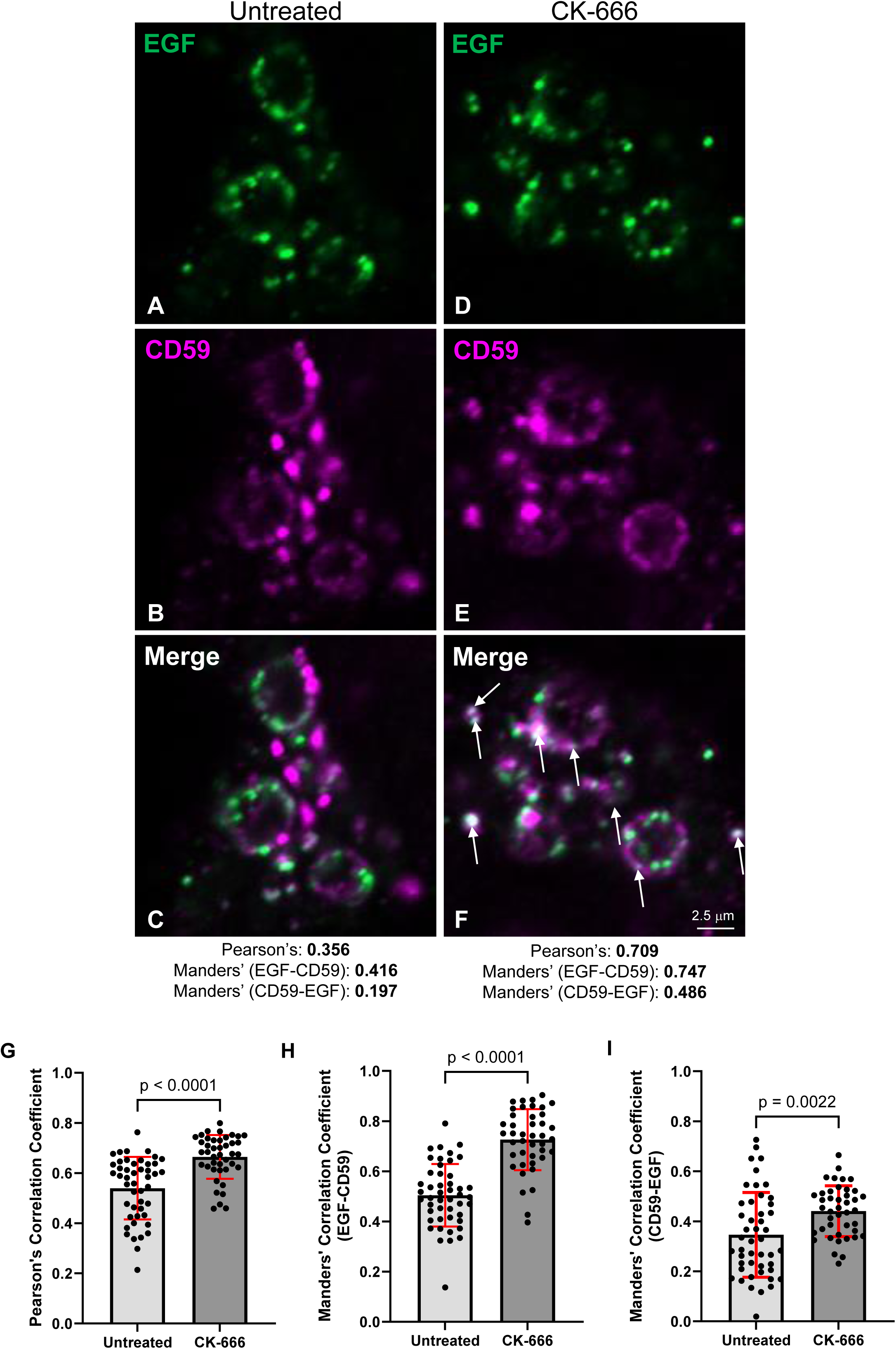
The degradative and retrieval subdomains on endosomes coalesce upon branched actin inhibition. **(A-F)**. HeLa cells were transfected with mCherry-RAB5 Q79L and were co-incubated with EGF-488 and anti-CD59 antibody for 20 min. During the last 5 min of uptake, either (**A-C**) no inhibitor or (**D-F**) 300 µM CK-666 was added to the media. Pearson’s and Manders’ correlation coefficients for each representative image are listed. **(G).** ImageJ was used to calculate Pearson’s correlation coefficient for 47 ROIs in the untreated group and 42 ROIs in the CK-666-treated group from three independent experiments. Statistical significance was determined using an unpaired two-tailed *t*-test. **(H, I)**. ImageJ was used to calculate Manders’ correlation coefficients (M1 and M2) for 47 ROIs in the untreated group and 42 ROIs in the CK-666-treated group from three independent experiments. Statistical significance was determined using an unpaired two-tailed *t*-test.

## DISCUSSION

Actin polymerization is required at multiple steps along endocytic trafficking pathways, including cargo internalization through diverse endocytic mechanisms at the PM and regulation of endosome dynamics (Chakrabarti *et al*, 2021). At the endosome, actin polymerization is required for endosome fission and receptor recycling, endosome motility, and the early-to-late endosome transition (Chakrabarti *et al*., 2021; Muriel *et al*, 2016). Formin-mediated linear actin polymerization is a mechanism for actin-based endosome motility (Gasman *et al*., 2003; Wallar *et al*., 2007). In contrast, the WASH complex promotes ARP2/3-mediated branched actin polymerization, which is required for endosome fission and receptor recycling (Derivery *et al*., 2012; Derivery *et al*., 2009; Gomez & Billadeau, 2009). Using a formin inhibitor and a specific ARP2/3 inhibitor, we validated the requirement for ARP2/3-mediated branched actin in endosome fission and receptor recycling, while demonstrating that formin-mediated linear actin is dispensable for these processes (Fig. 1 and 2). These findings highlight the presence of distinct pools of actin at the endosome, generated by different nucleation-promoting factors, that regulate separate endosomal functions.

To understand how ARP2/3-mediated actin supports endosome function, we next investigated its spatial relationship to cargo at the endosome membrane. The WASH complex is recruited to endosomes by the retromer (Gomez & Billadeau, 2009; Harbour *et al*., 2012; Jia *et al*., 2012), and the WASH, retromer, and retriever complexes localize to shared regions on endosomes, referred to as the retrieval domain (Derivery *et al*., 2012; McNally *et al*., 2017; Simonetti & Cullen, 2019). However, the spatial relationship between branched actin and specific cargos within these domains has not been studied. Addressing this question is challenging, because endosomes are small and the actin network at these structures is highly dynamic. To overcome this limitation, we utilized a constitutively active RAB5 mutant to enlarge endosomes and allow visualization of the spatial relationship between actin and select cargos. We found that Tf localizes to regions on the endosome that lie adjacent to regions of branched actin (Fig. 5). We refer to cargo domains adjacent to actin as “bounded,” and ∼60% of Tf-containing domains met this criterion under untreated conditions (Fig. 5).These observations reveal a spatial association between cargo domains and branched actin on the endosome, and support a model in which ARP2/3-mediated actin polymerization acts as a physical barrier that confines cargo within discrete endosomal membrane domains. Having established this spatial relationship, we next asked how branched actin might mechanistically confine cargo within these regions.

Inhibition of branched actin polymerization with an ARP2/3 inhibitor led to a more diffuse pattern of Tf and EGF around the endosome (Fig. 6 and 7), indicating that branched actin contributes to the confinement of cargo within discrete endosomal domains. The mechanism by which branched actin maintains this segregation remains unclear. At the PM, lateral diffusion of receptors is regulated by several factors, including the cytoskeleton, protein density, membrane curvature, and lipid domains (Fujiwara *et al*, 2016; Jacobson *et al*, 2019; Maynard & Triller, 2019). In contrast, the mechanisms regulating receptor diffusion at endosomes are not as well characterized. Several mechanisms may explain how actin contributes to cargo segregation within endosomal domains. One possibility is that cargos “bounded” by actin experience reduced lateral diffusion due to steric hindrance imposed by adjacent regions of branched actin (Fujiwara *et al*., 2016). Alternatively, direct interactions between receptors and the actin network may contribute to cargo confinement. For example, EGFR and membrane type 1–matrix metalloproteinase (MT1-MMP) contain intracellular actin-binding domains, suggesting that direct receptor-actin interactions could influence their spatial segregation on endosomes (den Hartigh *et al*, 1992; Yu *et al*, 2012). Consistent with this notion, a chimeric TfR containing an actin-binding domain was sufficient to sort TfR-Ub away from a degradative fate (MacDonald *et al*., 2018). A third possibility is that actin-mediated membrane remodeling contributes to cargo segregation. Diffusion of TfR into endosomal tubules occurs ∼4 times faster than diffusion of β2AR, highlighting receptor-specific diffusion dynamics as a potential sorting mechanism (Puthenveedu *et al*., 2010). Since actin stabilizes and promotes membrane tubulation (Puthenveedu *et al*., 2010; Simonetti & Cullen, 2019), actin-dependent tubule formation could facilitate cargo segregation by selectively promoting the entry of receptors with different diffusion properties into these structures.

Our data are consistent with ARP2/3-mediated branched actin confining cargo to discrete regions on the endosomal membrane, thereby helping to maintain established subdomains. The initial observation that RAB4, RAB5, and RAB11 localize to distinct regions on the same endosomes led to the proposal that endosomes are organized into spatially distinct subdomains (Sonnichsen *et al*., 2000). Since then, several mechanisms involved in the establishment of these subdomains have been identified, including SNX proteins, protein sorting complexes, and phosphoinositide conversion (McNally *et al*., 2017; Popoff *et al*., 2009; Posor *et al*, 2022; Temkin *et al*., 2011; Zhang *et al*., 2018). Here, we show that inhibition of ARP2/3 leads to coalescence of the degradative and retrieval subdomains (Fig. 8), indicating that branched actin polymerization represents an additional mechanism that contributes to endosomal subdomain maintenance. How protein sorting complexes, the actin network, and the lipid environment cooperate to establish and maintain these subdomains remains unclear. The phosphoinositide 3-kinase (PI3K) Vps34 is recruited to EEs by RAB5, where it generates PI3P (Christoforidis *et al*, 1999; Posor *et al*., 2022). Proteins involved in cargo sorting, including SNX17, SNX27, WASH complex, and other FYVE domain-containing proteins, are subsequently recruited and stabilized at the EE through direct interactions with PI3P (Chandra *et al*, 2019; Cullen & Steinberg, 2018; Jia *et al*, 2010; Rivero-Rios *et al*, 2025). These observations suggest that phosphoinositide conversion may establish a hierarchical framework that enables cargo sorting into distinct subdomains.

Interestingly, a pool of PI4P is also present at EEs and partially colocalizes with Tf, and inhibition of PtdIns4KIIα delays receptor recycling (Henmi *et al*, 2016). It has therefore been proposed that conversion of PI3P to PI4P within the retrieval domain of endosomes contributes to efficient receptor sorting and recycling (Dong *et al*, 2016; Henmi *et al*., 2016; Jani *et al*, 2022; Posor *et al*., 2022). Future studies will be required to determine how phosphoinositide conversion, actin polymerization, and cargo-sorting complexes are coordinated spatially and temporally to regulate the establishment and maintenance of endosomal subdomains.

Given the role of branched actin in subdomain maintenance, an important question is how disruption of actin impacts cargo trafficking outcomes. Inhibiting branched actin polymerization causes cargo accumulation in the EE (Fig. 3). However, since branched actin is required for both sorting and fission, it is unclear how defects in subdomain maintenance and cargo segregation influence trafficking to downstream compartments. Additional studies that uncouple cargo sorting and endosome fission will be required to determine how cargo mis-sorting alters trafficking outcomes. One approach may be to disrupt endosomal subdomain establishment through actin-independent mechanisms, such as inhibition of lipid kinases or disruption of SNX-cargo interactions.

While our study highlights a spatial relationship between receptors and branched actin at endosomes, it also raises questions about the temporal regulation of actin during endosomal trafficking. For example, are there actin regulatory proteins that participate in endosome fission but not receptor sorting, or vice versa? We previously showed that FCHSD2 promotes branched actin polymerization to enable endosome fission (Frisby *et al*., 2024), but its role in cargo sorting is unknown. Because FCHSD2 knockout significantly reduces branched actin at EEs (Frisby *et al*., 2024), it is possible that this protein also contributes to cargo sorting. Determining whether endosomal subdomains are disrupted in FCHSD2 knockout cells will therefore be important. In addition to actin polymerization, actin depolymerization is also required for fission and recycling (Dhawan *et al*., 2022; Noll *et al*, 2019), underscoring the importance of temporal control of actin dynamics. Coro2A, an actin regulatory protein at endosomes that negatively regulates actin polymerization (Dhawan *et al*., 2022), may play a role in this process. Our current model proposes that Coro2A promotes depolymerization during the terminal steps of fission, clearing steric obstruction and allowing EHD1 access to the neck of the tubulated endosome (Deo *et al*., 2018; Dhawan *et al*., 2022; Naslavsky & Caplan, 2023). It will therefore be important to determine whether Coro2A also functions earlier in cargo sorting and subdomain organization.

The actin network at the endosome is highly dynamic. Indeed, fluorescence recovery after photobleaching (FRAP) experiments have shown that endosome-associated actin recovers within ∼20 seconds (Puthenveedu *et al*., 2010). This rapid turnover underscores the transient nature of endosomal actin and suggests that continuous actin remodeling is required to coordinate the dynamic processes governing endosome organization and fission. Our findings highlight ARP2/3-mediated branched actin as a key regulator of cargo segregation, endosome subdomain maintenance, and endosome fission.

## MATERIALS AND METHODS

### Antibodies and reagents

The following antibodies were used: anti-CD59 (purified MEM-43/5, CLX64AP Cedarlane Labs, 1:200), anti-cortactin (05-180-I, Sigma, 1:200), Alexa Fluor 647-conjugated goat anti-mouse (115-606-008, Jackson ImmunoResearch, 1:750), Alexa Fluor 488-conjugated goat anti-rabbit (A11034, Molecular Probes, 1:500), and Alexa Fluor 568-conjugated goat anti-rabbit (A11036, Molecular Probes, 1:500). The mCherry-RAB5 Q79L (35138, Addgene) plasmid was used to generate enlarged endosomes. The following reagents were used: Alexa Fluor 488-conjugated transferrin (T13342, Invitrogen, 1:700), Alexa Fluor 488-conjugated EGF (E13345, Invitrogen, 1:100), CF-488-conjugated Phalloidin (42-T VWR, Biotium, 1:700), SMIFH2 (344092, MilliporeSigma, 25 µM), CK-689 (182517, MilliporeSigma, 300 µM), and CK-666 (182515, MilliporeSigma, 300 µM).

### Cell culture

HeLa cells (ATCC-CCL-2) were cultured with complete DMEM (high glucose) (ThermoFisher Scientific, Carlsbad, CA) containing 10% fetal bovine serum (FBS) (Sigma-Aldrich), 1× penicillin-streptomycin, 100 µg/mL Normicin, and 2 mM L-glutamine at 37°C in a humidified incubator with 5% CO₂. Transfection with mCherry-RAB5 Q79L (35138, Addgene) was achieved using the GeneExpresso transfection system (Excellgen, Rockville, MD). All transfections were performed overnight and used 1.5 µg of DNA for a 35 mm plate.

### Actin inhibitors

HeLa cells plated on coverslips were transfected with mCherry-RAB5 Q79L using the GeneExpresso transfection system. The following day, cells on coverslips were treated with SMIFH2, CK-689, or CK-666 for 20 min. Coverslips in the “untreated” condition had their medium replaced with complete DMEM and experiments were performed in parallel with the treated groups. For all experiments in this study, the formin inhibitor SMIFH2 was used at 25 µM, the inactive control CK-689 was used at 300 µM, and the ARP2/3 inhibitor CK-666 was used at 300 µM. Following 20 min of treatment, the cells were washed with PBS and fixed with 4% paraformaldehyde diluted in PBS for 20 min. After fixation, the coverslips were washed with PBS three times and immunostained for 1 h with anti-cortactin, to mark branched actin, and 488-phalloidin, to mark all filamentous actin. Cells were then washed three times with PBS and incubated for 30 min with the appropriate 647 fluorochrome–conjugated secondary antibody. Confocal images were captured using a Zeiss LSM 800 confocal microscope (CarlZeiss) with a 63 ×/1.4 NA oil objective.

The total number of mCherry-RAB5 Q79L endosomes was counted per image. The percentage of the RAB5Q79L endosomes that contact cortactin was expressed as a percentage of the total number of enlarged endosomes for each image. This quantification was also done for RAB5Q79L endosomes that contact phalloidin.

### Measurement of endosome size

HeLa cells were plated on coverslips and incubated with complete medium or the respective inhibitor diluted in complete media (see “Actin Inhibitors” for dilutions). Coverslips were washed and fixed with 4% paraformaldehyde at 15 min, 30 min, and 50 min time points. Following fixation, coverslips were washed three times with PBS and co-stained with primary antibodies against EEA1 and cortactin. Coverslips were washed with PBS three more times and stained with the appropriate 488-and 568-conjugated secondary antibodies. After the coverslips were mounted, images were captured using a Zeiss LSM 800 confocal microscope (CarlZeiss) with a 63 ×/1.4 NA oil objective.

The size of EEA1-decorated endosomes was measured using Imaris 9.9.1 (Oxford Instruments, Abingdon, United Kingdom) by creating surfaces with the parameters found in Table S1, row 1. Endosome area was exported from Imaris, and the calculated areas for each treatment group were normalized to the mean of the untreated group at each time point and plotted. The total number of endosomes analyzed was ∼50,000 per treatment group from three independent experiments.

### Tf recycling assay

For the transferrin recycling assay, HeLa cells were incubated with Transferrin-AlexaFluor 488 (Tf-488; Invitrogen) diluted 1:700 in complete DMEM. SMIFH2, CK-689, CK-666, or no inhibitor was added to the Tf-containing medium (dilutions listed above). Cells on coverslips were incubated with Tf-488 and the specified inhibitor for 10 minutes of uptake. After uptake, cells were either washed with PBS and fixed, or cells were washed with PBS and chased for 40 min with complete media containing the inhibitors. Following complete media chase, cells were washed with PBS, fixed with 4% paraformaldehyde for 20 min, and mounted. Images were captured using a Zeiss LSM 800 confocal microscope (CarlZeiss) with a 63 ×/1.4 NA oil objective.

Using Zen Blue software, the arithmetic mean intensity for each uptake image was analyzed. The arithmetic mean intensities of the SMIFH2-, CK-689-, and CK-666-treated groups were normalized to the mean of the untreated group (set to 100%). To quantify “% of Tf remaining in cells” after chase, the arithmetic mean intensities of all the chase images were individually measured as a percentage of the average intensity of uptake (for that treatment) and plotted.

Three independent experiments were performed and quantified (30 images per treatment).

### EEA1 cargo localization

HeLa cells were incubated with Tf-488 for 10 min to allow cargo uptake. Following uptake, coverslips were washed once with PBS and either fixed immediately or chased in complete medium containing no inhibitor or CK-666 (300 µM). Cells were fixed with 4% paraformaldehyde at 15, 30, or 45 min during the chase. Following fixation, coverslips were washed with PBS and incubated with a primary antibody against EEA1, followed by three PBS washes and incubation with the appropriate secondary antibody. Coverslips were mounted and images were acquired using a Zeiss LSM 800 confocal microscope (Carl Zeiss) with a 63×/1.4 NA oil immersion objective.

Eight confocal images per experiment were collected across three independent experiments (24 total images) and quantified using Imaris 9.9.1 (Oxford Instruments). Tf-containing vesicles (Table S1, row 2) and EEA1-decorated endosomes (Table S1, row 4) were three-dimensionally rendered in Imaris using the Surfaces function with the specified parameters. Using the “Detailed Statistics” module, the total volume of Tf-positive structures and the volume of Tf overlapping with EEA1 were exported. The volumetric sum of Tf overlapping with EEA1 was expressed as a percentage of the total Tf volume for each image, yielding the percentage of Tf localized to EEA1-positive endosomes. This analysis was performed for 24 images for each time point (0, 15, 30, and 45 min).

EGF localization to EEA1-positive endosomes was measured using the same procedure, except that cells were serum-starved for 1 h prior to the 10 min uptake period. EGF-containing vesicles and EEA1-positive endosomes were rendered using the respective Imaris parameters (Table S1, row 3). The total EGF volume and the volume overlapping with EEA1 were exported, and the percentage of EGF localized to EEA1-positive endosomes was calculated for 24 images per time point (0, 15, 30, and 45 min).

### Quantification of cortactin bounding Tf

HeLa cells were transfected with mCherry-RAB5 Q79L using the GeneExpresso transfection system overnight. The next day, cells were incubated with no inhibitor, SMIFH2, CK-689, or CK-666 diluted in complete DMEM (see “Actin inhibitors” for dilutions) for 4 min. Following this pre-treatment, Tf-488 was added to the inhibitor-containing media for 6 min of uptake. Cells were then washed with PBS, fixed with 4% paraformaldehyde, and stained with a primary antibody against cortactin and the appropriate secondary antibody. Following staining, coverslips were washed with PBS three times and mounted. A Zeiss LSM 800 confocal microscope (Carl Zeiss) with a 63 ×/1.4 NA oil objective was used to capture images.

The regions of Tf were analyzed in ImageJ (National Institutes of Health, Bethesda, MD). The circle tool was used to draw a circle along the mCherry-RAB5 Q79L endosome membranes. The drawn circle was added to the ROI manager so that the coordinates for the circle remained the same for the image in each channel (488, 568, and 647). The workflow for each channel after drawing the circle is as follows: plugins>new>macro>(pasted the code for a macro we wrote)>run. The code for the macro used was deposited at Zenodo (<u>10.5281/zenodo.18893507</u>).

The script for the macro calculated the fluorescence intensity at each degree along the drawn circle, allowing us to develop a fluorescence intensity profile of a circle. Each fluorescence intensity profile is representative of one endosome. In Microsoft Excel (Redmond, WA), the raw fluorescence intensity at each degree was normalized as a percentage of the mean for each channel independently.

To quantify Tf “bounded” by cortactin, the normalized fluorescence intensities for Tf and cortactin were analyzed in RStudio (Posit PBC, Boston, MA). The RStudio packages that were installed included *readxl* (v.1.4.5)(Wickham, 2025), *openxlsx* (v4.2.8)(Schauberger, 2025), and *pracma* (v2.4.4)(Borchers, 2023). First, “*findpeaks*” was used to mark all of the maximal points of Tf (peak defined as: “maxima in a time series). Then, a code was written to identify all peaks of Tf that had a maximal fluorescence intensity greater than 130% (code deposited at Zenodo: (10.5281/zenodo.18893540). A Tf peak was considered “bounded” if the peak occurred within ± 20 degrees of a cortactin peak above 130%, and these “bounded” peaks were labeled with a red dot. If the Tf peak did not contain a cortactin peak above 130% ± 20 degrees, it was considered “not bounded” and was labeled with a gray dot. The script then calculated the percentage of Tf peaks per endosome that were bounded by cortactin vs. the total Tf peaks on that endosome.

From three independent experiments, 72 Tf peaks from 37 endosomes for the control group, 70 peaks from 43 endosomes for SMIFH2-treated, 73 peaks from 38 endosomes for 689-treated, and 70 peaks from 37 endosomes were quantified.

### Quantification of Tf and EGF domain width on endosomes

To measure the width of Tf-containing domains, cells were transfected with mCherry-RAB5 Q79L using the GeneExpresso system, and the same Tf uptake methods were used as described in “*Quantification of cortactin bounding Tf”*. Fluorescence intensity profiles along endosome membranes were extracted in ImageJ and normalized in Microsoft Excel as described above. For each Tf-containing domain, the maximal fluorescence intensity was identified and defined as the center of the peak. The maximal fluorescence intensity for each peak was set to 100%, and fluorescence intensities at each degree within the ± 20 degrees window were normalized to this maximal value (all 41 normalized degrees comprise a Tf “peak”).

The plotted curves represent the mean normalized fluorescence intensity at each degree for all Tf peaks within each treatment group across three independent experiments (control: 72 Tf peaks from 37 endosomes; SMIFH2-treated: 70 peaks from 43 endosomes; CK-689-treated: 73 peaks from 38 endosomes; CK-666-treated: 70 peaks from 37 endosomes).

EGF experiments were performed using the same 4 min pretreatment protocol, except that EGF-488 was added for a 17 min uptake period. Quantification of EGF peaks was performed identically to the Tf peak analysis described above. Across three independent experiments, the following peaks were analyzed: control (84 peaks from 47 endosomes), SMIFH2-treated (86 peaks from 51 endosomes), CK-689-treated (72 peaks from 39 endosomes), and CK-666-treated (86 peaks from 49 endosomes).

### EGF and CD59 co-uptake

HeLa cells were transfected with mCherry-RAB5 Q79L using the GeneExpresso transfection system. The following day, cells plated on coverslips were incubated with Alexa Fluor 488-conjugated EGF (EGF-488) and an anti-CD59 antibody diluted in complete medium for 20 min to allow co-uptake. During the final 5 min of uptake (beginning at the 15 min time point), CK-666 (300 µM) or no inhibitor was added to the medium.

Following uptake, cells were incubated with an acid stripping solution (0.5 M NaCl, 0.5% acetic acid, pH 3.0) for 2 min to remove antibodies remaining on the cell surface. Cells were washed twice with PBS, fixed with 4% paraformaldehyde for 20 min, and then incubated with the appropriate Alexa Fluor 647-conjugated secondary antibody to label internalized CD59.

Coverslips were mounted and images were acquired using a Zeiss LSM 800 confocal microscope (Carl Zeiss) with a 63×/1.4 NA oil immersion objective.

Colocalization of EGF and CD59 within endosomes was analyzed using ImageJ (National Institutes of Health). Regions of interest (ROIs) corresponding to mCherry-RAB5 Q79L-positive endosomes were defined for each image. Colocalization coefficients were calculated using the Just Another Colocalization Plugin (JACoP; v2.1.1)(Bolte & Cordelieres, 2006). Pearson’s correlation coefficient and Manders’ coefficients (M1 and M2) were calculated for each ROI. Three independent experiments were performed. Colocalization values were plotted for 47 ROIs in the untreated group and 42 ROIs in the CK-666-treated group.

### Statistics

Data were collected from three independent experiments unless otherwise indicated. Mean values ± standard deviation (SD) were plotted for each dataset. Graphs and statistical analyses were generated using GraphPad Prism (v10.2.3).

Normality was assessed using the D’Agostino-Pearson or Shapiro-Wilk tests. For comparisons between two groups, an unpaired two-tailed Student’s *t*-test was used when data met the assumption of normality. If normality was not met, a two-tailed Mann-Whitney nonparametric test was used. Statistical significance was defined as *p* < 0.05.

## DATA AVAILABILITY

The following custom-written code was deposited in Zenodo:

1) Circular ROI code (ImageJ) 10.5281/zenodo.18893507
2) Peak code (RStudio) 10.5281/zenodo.18893540

---

**Table S1.** Imaris parameters used to render surfaces.

| **Reference number** | **Description** | **Surface grain size (μm)** | **Diameter of largest sphere (μm)** | **Manual threshold value** | **Region growing estimated diameter (μm)** | **Number of Voxels above** |
| --- | --- | --- | --- | --- | --- | --- |
| **1** | **Measurement of endosome size:**<br>EEA1 | 0.198 | 0.200 | 753.304 | 0.990 | N/A |
| **2** | **EEA1 cargo localization:**<br>Transferrin structures | 0.198 | 0.300 | 707.243 | N/A | 28.6 |
| **3** | **EEA1 cargo localization:**<br>EGF structures | 0.198 | 0.300 | 2079.55 | N/A | N/A |
| **4** | **EEA1 cargo localization:**<br>EEA1 | 0.198 | 0.300 | 1612.23 | N/A | N/A |

